# The genome and metabolome of the tobacco tree, *Nicotiana glauca*: a potential renewable feedstock for the bioeconomy

**DOI:** 10.1101/351429

**Authors:** Björn Usade, Takayuki Tohge, Federico Scossa, Nicolas Sierro, Maximilian Schmidt, Alexander Vogel, Anthony Bolger, Amanda Kozlo, Eugenia MA Enfissi, Kris Morrel, Manuel Regenauer, Asis Hallab, Colin Ruprecht, Heidrun Gundlach, Manuel Spannagl, Yaw Koram, Klaus FX Mayer, Wout Boerjan, Paul D. Fraser, Staffan Persson, Nikolai V. Ivanov, Alisdair R. Fernie

## Abstract

**Background:** Given its tolerance to stress and its richness in particular secondary metabolites, the tobacco tree, *Nicotiana glauca*, has been considered a promising biorefinery feedstock that would not be competitive with food and fodder crops.

**Results:** Here we present a 3.5 Gbp draft sequence and annotation of the genome of *N. glauca* spanning 731,465 scaffold sequences, with an N50 size of approximately 92 kbases. Furthermore, we supply a comprehensive transcriptome and metabolome analysis of leaf development comprising multiple techniques and platforms.

The genome sequence is predicted to cover nearly 80% of the estimated total genome size of *N. glauca*. With 73,799 genes predicted and a BUSCO score of 94.9%, we have assembled the majority of gene-rich regions successfully. RNA-Seq data revealed stage-and/or tissue-specific expression of genes, and we determined a general trend of a decrease of tricarboxylic acid cycle metabolites and an increase of terpenoids as well as some of their corresponding transcripts during leaf development.

**Conclusion:** The *N. glauca* draft genome and its detailed transcriptome, together with paired metabolite data, constitute a resource for future studies of valuable compound analysis in tobacco species and present the first steps towards a further resolution of phylogenetic, whole genome studies in tobacco.

## Background

Tree tobacco (*Nicotiana glauca*) is a wild tobacco species, not only endemic to South America but also widespread as an introduced, invasive species in other continents, that exhibits partial drought and salinity tolerance under high irradiance [1]. *N. glauca* grows in arid environments [1], usually not suitable for food crops, and has vast potential as a biofuel stock due to an exceptionally high content of hydrocarbons [2]. In addition, *N. glauca* has been discussed for medicinal purposes due to its rich content in metabolites such as scopoletin [3]. The genome size of the diploid (2n=24) *N. glauca* was estimated previously at 5.33pg (5.2 GB) using Feulgen staining [4, 5], which, surprisingly, is larger than the cultivated, allotetraploid *N. tabacum (2n=4×=48) [6]. N. glauca* is additionally reported to be a source of vitamin D [7]; however, these data need to be substantiated.

While *N. glauca* is usually counted towards the *Nicotiana* section *Noctiflorae*, which is also supported by RNA-Seq data [8], there are indications for it having resulted from the hybridization between a member of the section *Noctiflorae* and one from the section *Petunioides* [9]. Such hybridization events have been used to explain the invasiveness of species [10]. The provenience of *N. glauca* should be discernable given proper genomic data. While recent years have seen an avalanche of plant genomes [11, 12], there is comparatively little data on this interesting tobacco clade, and any significant efforts made have been primarily targeting domesticated tobacco and its likely progenitors [6, 13, 14], thus not offering insights into the origin of *N. glauca*. The only exception that might shed light on the origin of *N. glauca* is the genome of *N. attenuata* (Section *Petunioides*), which was used to study the evolution of nicotine biosynthesis [15]. However, there is no genome sequence of an unequivocal member of the section *Noctiflorae*, which would be necessary to classify *N. glauca*.

Beyond these basic biological questions, *N. glauca* is quite interesting from an applied perspective. This relates to the ever-increasing world population and the corresponding increase in energy requirements, coupled to environmental deterioration, which has become known as the food, energy, and environment trilemma [16]. Thus, development of novel biofuel or biomaterial feedstocks that are less detrimental than fossil fuels and do not compete for arable land with food and feed production are desired. Given the growth requirements and the amounts of hydrocarbon production of *N. glauca*, it would be ideally suited in the context of such new feedstocks.

Here, we report the genome, transcriptome, and comprehensive leaf metabolome of *N. glauca* in a growth cycle and provide case study field trials of the crop in arid environments. Given the rich complement of metabolites extractable from *N. glauca* and its yield data in arid environments, our work highlights a route for biofuel and/or biomaterial production in a manner that is noncompetitive to food production.

## Data Description

We extracted genomic nuclear-enriched DNA from young leaves of two *N. glauca* cultivars, following established protocols [6, 17]. Subsequently, we sequenced the DNA using an lllumina HiSeq for paired-end sequencing and an Illumina MiSeq for mate pair sequencing. We generated three libraries each of 4k and 8k as well as six libraries of 10 kbases or larger. The genome was assembled using SOAPdenovo2 [18] after removing clonal mate pair artifacts yielding a genome size of 3.51 Gbp and an N50 value of 91 kbases.

We isolated RNA from leaves at 10 different developmental stages as well as from stems, roots, and flowers, as described previously [19], and subsequently sequenced them. Metabolites were extracted from leaves, as described previously [19], and analyzed on different platforms.

## Analysis

### Genome sequencing and assembly

We sequenced the genome of *N. glauca* using paired-end data with insert sizes ranging between 200 and 730 bases. In total, we generated about 380 Gbases of sequence with a length of 101 base pairs. After adapter trimming and removing low-quality bases, 340 Gbases were available for assembly. Furthermore, we generated approximately 44, 78, and 26 million mate pair reads with insert sizes of approximately 4, 8, and 11-16 kbases, respectively.

We also survey-sequenced a second *N. glauca* cultivar (TW53) using 150 GB paired-end and small-insert data only. Based on a k-mer analysis of the genome size, we estimated the genome to be in the range of 4.5 Gbases for the first accession and of approximately 4.3 Gbases for *N. glauca* TW53. While the estimated genome sizes for both cultivars were quite consistent, they are somewhat smaller than the experimentally derived value using Feulgen staining of approximately 5.2 Gbases [4, 5] but within typical error margins.

We assembled the genome of the first cultivar using SOAPdenovo2 [18] with a k-mer size of 75 and obtained a scaffold N50 of 92 kbases. This is smaller than the N50s obtained for the domesticated tobacco of approximately 350 kbases [6] and much smaller than the N50 for the *N. tabacum* genome scaffolded by optical mapping [14]. However, its genome coverage is in line with the similar genome size of *N. tabacum*, where also approximately 3.5 Gbases (approximately 78%) of the genome were assembled. The remainder is likely comprised of “difficult” repetitive elements that are not easily resolvable using short-read sequencing technology, as is frequent for plant genomes [11, 20]. In order to assess the gene completeness of the genome, we used BUSCO [21] and our RNA-Seq data for *N. glauca*. For BUSCO, we obtained a gene completeness rate of 94.9% (Table 1), and we observed RNA-Seq mapping rates of approximately 82-92% of the reads using standard mapping parameters with salmon [22], Taken together, these data indicate that the genome assembly covers the coding capacity of *N. glauca* well.

**Table 1.**
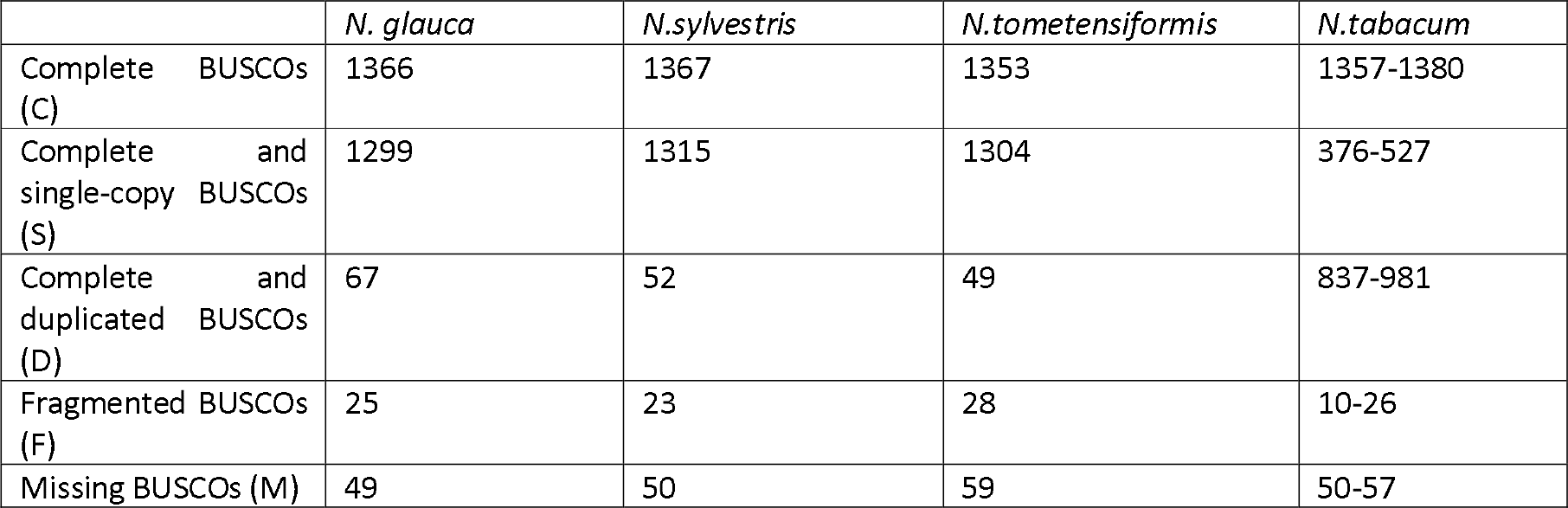
BUSCO analysis for gene completeness for *N. glauca*, *N. sylvestris*, *N. tometensiformis* and *N.tabacum*.

### Genome annotation

We performed gene annotation using AUGUSTUS, relying on more than 100 Gbases of RNA-Seq data in more than one billion reads covering different tissues, as well as statistical models of genes. In total, this led to the identification of 73,799 genes, which is in line with the nearly 100k genes typically found in tobacco species of similar genome size [6, 13–15].

We functionally annotated the gene models using the automatic Mercator plant genome annotation pipeline [23, 24] and compared them with other tobacco species. We also compared them with the high-quality genome of the reference species tomato [25] as well as with other Solanaceous species using the same pipeline. These analyses support the BUSCO data, as it was possible to find genes for almost all classes in *N. glauca*. To provide a more quantitative visual assessment, we counted the resulting annotations and displayed the annotations as sunburst charts. As Mercator uses a redundancy-reduced hierarchy, this gives a good indication of the covered gene functions and of the percentage of genes assigned to different categories. Indeed, in line with the gene analysis, this showed that the representation of different gene functions is highly conserved within the Solanaceae (Figure 1).

**Figure 1.**
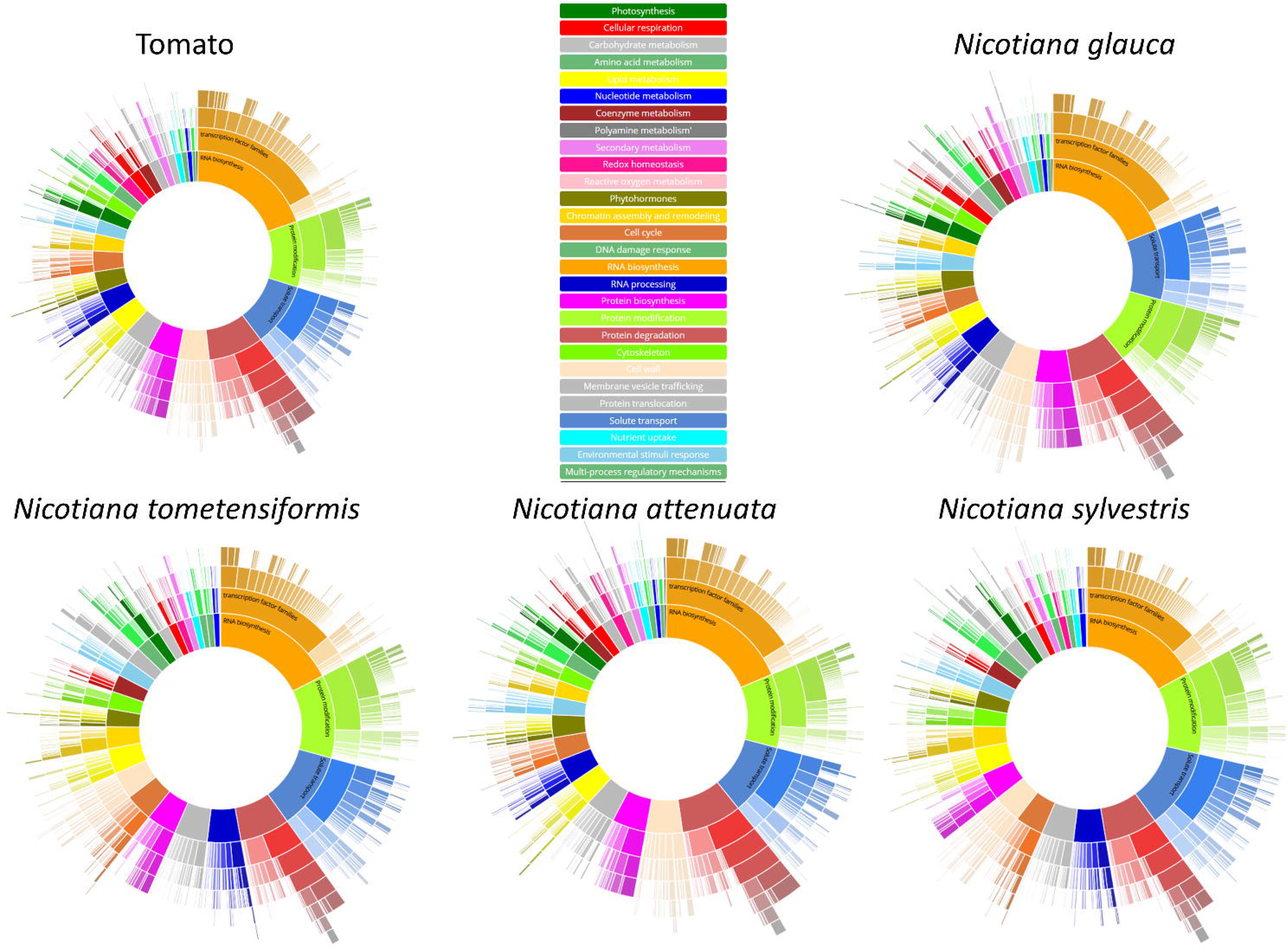
Gene function analysis of *N. glauca*, *N. attenuata*, *N. sylvestris*, and *N. tomentosiformis* as well as the Solanaceous reference tomato. For data analysis, we submitted the protein-coding genes or cDNA sequences to Mercator version X4.03, counted the resulting annotations, and summarized them as a sunburn chart, where the inner ring represents broad categories and further outer rings represent more refined subcategories. Data are available as an interactive visualization in the supplement, as well.

### Repetitive element analysis

Analysis of genomic repetitive elements revealed that most of them were Class I-type-derived. These were comprised primarily of long terminal repeat elements, where we found more Gypsy than Copia elements. However, we identified fewer repetitive elements than expected, in line with the large remaining gaps in the genome.

### Gene expression analysis through a developmental time course

To analyze leaf growth, we took samples through the development. To this aim, we harvested leaf material in duplicate at 10 different stages determined by leaf size (Supplemental Figure 1) and subjected them to RNA-Seq-based data analysis using salmon [22], As shown in Supplemental Figure 2, the replicates grouped together, and we observed a general separation by developmental time when data was subjected to multidimensional scaling. We confirmed this grouping by adding duplicate data from flower, root, and stem samples (Supplemental Figure 2). In total, more than 100 Gbases represented by 1.1 billion reads remained after trimming. Using these data, we analyzed differential gene expression in the leaf developmental time series using RobiNA [26] and edgeR [27] and identified 6,223 genes, which changed during the developmental time series (false discovery rate [FDR] p-value ≤0.05). We scaled and centered the differentially expressed genes and subjected them to k-means clustering. For this purpose, both silhouette [28] and elbow [29] methods showed that we obtained an optimal clustering by dividing the profiles into two clusters. Indeed, the predominant trend was either an upregulation of genes during the time course (cluster 1: 3,350 genes) or a continuous downregulation (cluster 2: 2,873 genes) (Figure 2).

**Figure 2.**
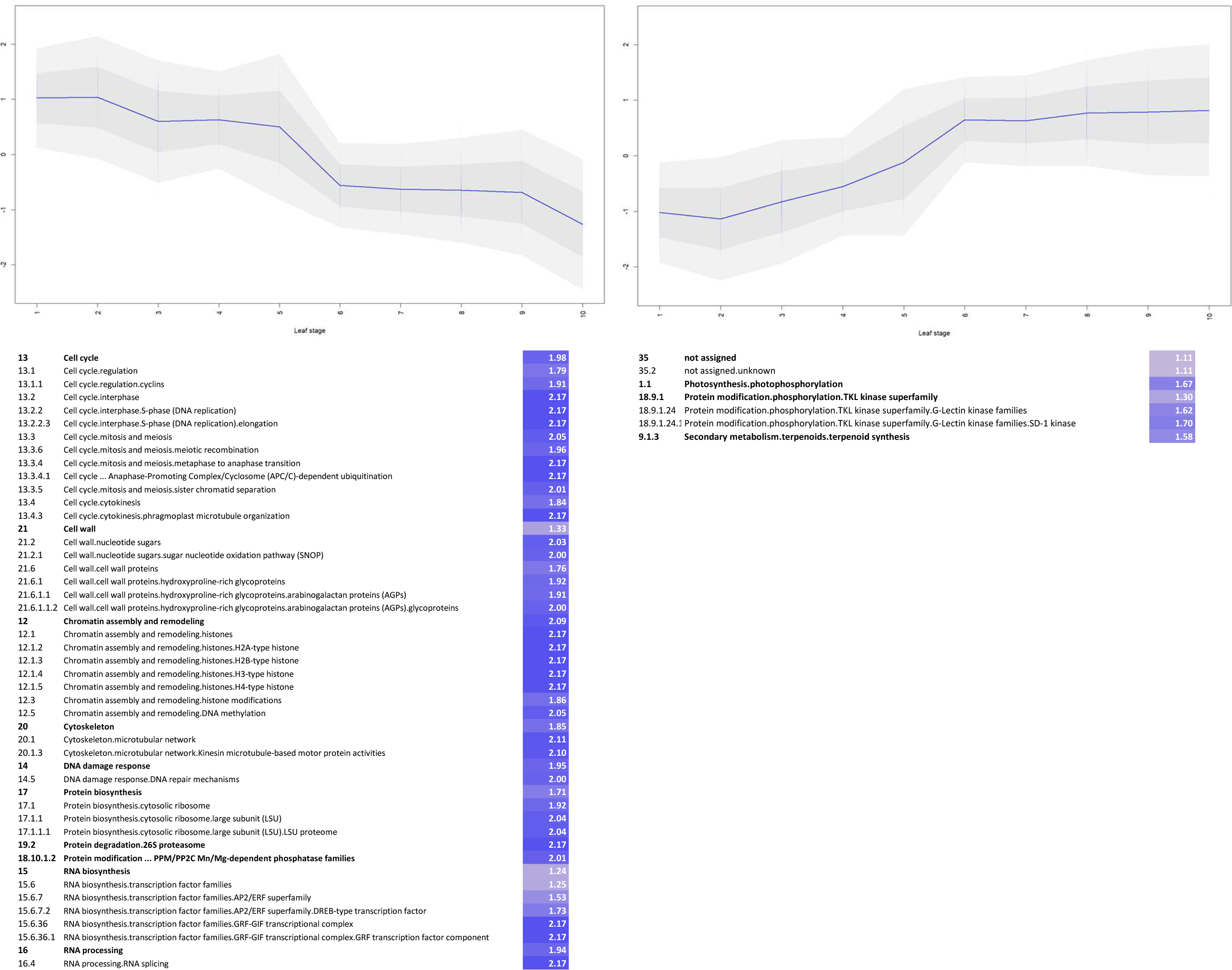
Visualization of the RNA-Seq developmental time course. For the developmental leaf time course, genes that were differentially expressed were extracted and genes were clustered using k-means clustering. Functional categories overrepresented in the clusters were determined using Fisher’s exact test, and overrepresented categories along with their enrichment versus all differentially expressed genes are shown.

Using Mercator classification and functional enrichment testing of each cluster versus the whole set of differentially expressed genes, we identified (i) photophosphorylation, (ii) tyrosine kinase-like (TKL) kinases, especially the SD1 type of TKL kinases, and (iii) terpenoid synthesis amongst the genes significantly enriched in the cluster representing upregulated genes (FDR≤0.05) (Figure 2).

In the cluster representing downregulated genes, we identified, as overrepresented categories, cell cycle, cytoskeleton, chromatin assembly, cell wall, and protein biosynthesis. These lists are in close agreement with previous plant leaf development time course data sets [30]. In addition, we found that the categories of DNA damage response, RNA biosynthesis and degradation, and 26S proteasome were overrepresented in this cluster (Figure 2).

### Metabolic profiles

Using three largely complementary and established metabolite profiling platforms, we also sampled greenhouse-grown *N. glauca* material during the 10 different stages of leaf development determined by leaf size (see above) using six replicates instead of two. After we centered and scaled the metabolite data, a principal component analysis revealed that PC1 (explaining 54% variance) discriminated the first five stages from the remaining stages (6-10) of leaf development and ordered almost all stages according to developmental time points, with the only exception of stages seven and eight, which changed their respective places. PC2 (explaining approximately 16% variance) provided further separation within the two main clusters from PC1. Given the separation by developmental stage, we considered the data of good quality and analyzed them further (Supplemental Figure 3).

After detailed analyses of the metabolite data (Supplemental Data), we observed an increase in lipids and terpenoid-related metabolites as well as in phenylpropanoids and related compounds, such as phenolamide, as the leaves developed and became older. Interestingly, the increase in terpenoid-related metabolites recapitulates the preferential enrichment of the corresponding genes in the upregulated gene cluster (Figure 2, Supplemental Figure 4). In the case of the phenylpropanoids, there was no concomitant enrichment. Indeed, we could only identify one PAL gene (*g66340*) whose expression was tagged as significantly different. However, this gene showed a rather inconsistent fluctuation, and a second gene (*g49941*) encoding a potential proteolytic PAL regulator was upregulated during the developmental time course. However, it is unlikely that these two genes would suffice to explain the observed trend of phenylpropanoid accumulation.

On the other hand, metabolites of the tricarboxylic acid cycle and polyamines were generally decreasing over the developmental stages (Supplemental Figure 4, Supplemental Data). While P-sitosterol increased only slightly during development, carotene increased more than five-fold. Conversely, quinate decreased approximately 60-fold, and citrate decreased approximately 30-fold. In addition, neither squalene nor some medium-to long-chain fatty acid (derivatives) were detectable in young leaf stages, but they accumulated considerably at later stages. Taken together, these data show that there is extensive developmental variation in the metabolic content of *N. glauca*. Hence, certain metabolites were enriched depending on the harvest time point. A glycoside of scopoletin, which has potential medicinal applications, varied substantially during development but peaked at stage 7, which thus could serve as a suitable harvest time to enrich for this metabolite.

Previously, vitamin D_3_ derivatives were identified in *N. glauca* leaves, and a content of approximately 1 ng/g fresh weight (FW) was estimated using a sensitive radioisotope assay [7], We therefore analyzed *N. glauca* plant material for the occurrence of vitamin D_3_; however, it remained below the detection limit, potentially due to lack of strong UV radiation [31].

### Analysis of cuticle biosynthesis genes

Because leaves of *N. glauca* have a thick, waxy layer and accumulate long-chain alkanes [2], which might improve drought tolerance, we analyzed genes related to cutin, suberin, and waxes, including both the biosynthesis of cutin and wax monomers and their export and assembly in the cuticular layer. For each step in the cuticle biosynthesis, we retrieved the sequences of the related enzymes from *N. glauca*, *N. tomentosiformis*, and *N. sylvestris* and compared their copy numbers, polymorphisms, and expression specificity across different tissues. We rated amino acid substitutions based on multivariate analysis of protein polymorphisms (MAPP) predictions, which estimate the amount of physicochemical violation of each nonsynonymous site in an alignment [32].

#### Cuticular lipid formation

The synthesis of cutin monomers is initiated with the hydroxylation of fatty acids in the endoplasmic reticulum of epidermal cells. The first hydroxylation is usually an ω-hydroxylation catalyzed by a cytochrome P450 of the CYP86 subfamily [33] (Figure 3). *N. glauca* harbors the three predicted CYP86A genes (*NGLAgSO*, *NGLAg31431*, *NGLA72346*), whereas *N. sylvestris* and *N. tomentosiformis* have only two predicted fatty acyl ω-hydroxylases. *NGLAgSO* is mostly expressed in roots, stems, and flowers, with lower expression during leaf development, while *NGLAg31431*, which seems to have no paralog in the other two species, and *NGLA72346* are highly expressed in leaf, stems, roots, and flowers.

**Figure 3.**
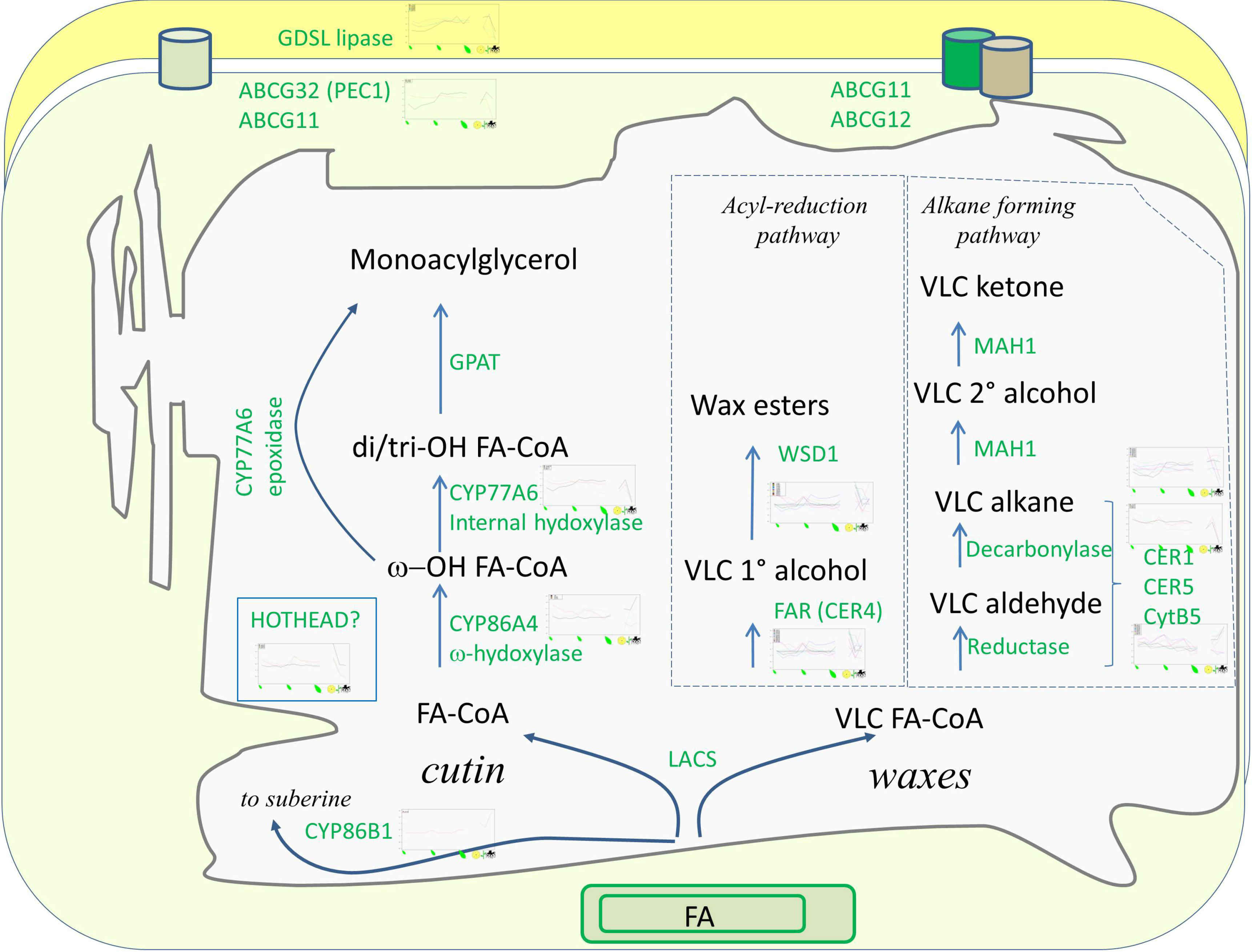
MapMan visualization of Cutin Biosynthesis. RNA-Seq data normalized to the average log expression, with data visualized on an overview scheme for cutin biosynthesis.

The next step in the synthesis of cutin monomers is the internal hydroxylation of the fatty acid chain catalyzed by a midchain hydroxylase of the CYP77 subfamily, allowing formation of ester linkages between simple monomers [34], *N. glauca*, *N. sylvestris*, and *N. tomentosiformis* have three CYP77 genes annotated as fatty acyl in-chain hydroxylases (two CYP77A1 and one CYP77A2). All *N. glauca* CYP77 genes are predominantly expressed in vegetative tissues, with lower expression in the roots.

Very long-chain fatty acids (VLCFA), mainly C22 and C24, are components of suberin, which is a lipid polymer present both in roots and in the aerial parts of the plant. Similar to cutin, suberin is composed of aliphatic monomers that undergo polymerization through the formation of ester linkages involving the carboxy terminus and the in-chain and co-hydroxy groups. The cytochrome genes involved in the ω-oxygenation of these VLCFAs belong to the CYP86B1 subfamily [35]. A single gene of the CYP86B1 subfamily is annotated as a very long-chain fatty acyl omega-hydroxylase in *N. glauca, N. sylvestris*, and *N. tomentosiformis*. This gene is predominantly expressed in roots and flowers.

Members of the GDSL esterase/lipase gene class catalyze the formation of higher-order cutin polymers, starting from dihydroxyglycerol esters of fatty acids. In *Arabidopsis* and in the three tobacco species, the family is composed of five members. The five genes in *N. glauca* are expressed ubiquitously, although some members show high expression in specific tissues. For example, *g64386* and *g66316* are highly expressed in stems, flowers, and leaves (compared with roots), while the level of *g13725*, which is expressed in flowers and stems, also increases during leaf development. The analysis of polymorphisms between *N. glauca* and other tobacco species highlighted several changes predicted to be intolerant. Among these, the *N. glauca* gene *g64386* has an amino acid substitution in a highly conserved region, where alanine is substituted by threonine; the change is predicted to impact protein function by MAPP (p-value=6.498E-4).

Another gene that might play a role in cutin biosynthesis is HOTHEAD, whose *Arabidopsis* mutants have a peculiar cutin composition, with a marked increase in co-hydroxy fatty acids. *N. glauca* has three ubiquitously expressed genes encoding HOTHEAD proteins, in contrast with *N. tomentosiformis* and *N. sylvestris*, which feature only two genes. Analysis of polymorphisms revealed several *N. glauca-*specific variations in otherwise conserved regions. Although these changes were not predicted to play a major role for protein function, conservative amino acid changes in other genes active in the cutin pathway have been shown to alter substrate specificity [36].

#### Acyl-reduction pathway

The reduction of VLCFA-coenzyme A (CoA) (the products of the fatty acid elongase complex) to primary alcohols is the first committed step in the synthesis of wax components. The genes catalyzing this step belong to the class of fatty acyl-CoA reductases (CER4 in *Arabidopsis* [37]). We identified 10 members in *N. glauca*. Some of the *N. glauca* genes show marked tissue specificity: *g17924* is expressed only in flowers, while *g29510* is a root-specific gene. Six acyl-CoA reductase genes are instead expressed ubiquitously across all tissues (*gl404*, *gl2813*, *gl5504*, *gl9042*, *g29192*, *g47356*). However, for *g47356*, we detected a positive selection signal. In phylogenetic tests [38–40], the gene and several conserved amino acids showed signs of positive selection. One of the selected amino acids falls within the Male sterility (MS), NAD-binding protein domain (IPR013120). The domain’s function is assumed to be essential for the formation of pollen wall substances and is a functional part of the fatty acyl reductase MS2 in *Arabidopsis* [41]. For the remaining two genes (*g29406* and *g40539*), we did not detect expression in any of the tissues.

MAPP identified several polymorphisms predicted to impact protein function. For example, the *N. glauca* gene *g1404* has two intolerant changes: (i) a substitution with Ser at position 253 (where all remaining sequences have either Lys, Gln, or Arg), and (ii) a substitution with Gln at position 574 (where all remaining sequences have either Asp, Glu, Lys, or Asn). Similarly, the gene *g19042* also has two intolerant changes: (i) a substitution with Val in position 237 (which is otherwise occupied by Phe, Ile, Leu, or Met in the multialignment), and (ii) a Leu substitution in position 353, located in an extremely conserved position where all remaining sequences instead have Phe. These changes may therefore indicate functional divergence of some members of acyl-CoA reductases in *N. glauca*.

##### Wax ester synthase and diacylglycerol acyltransferase (WSD)

Primary alcohols, the products of acyl-CoA reductases, can be esterified to C16/C18 fatty acyl-CoA by the action of a wax ester synthase represented by WSD1 in *Arabidopsis*. The annotation by Mercator showed the presence of eight wax ester synthase genes in *N. glauca*. This class thus represents an expansion with respect to *N. sylvestris* and *N. tomentosiformis*, for which we could only identify five and two wax ester synthase genes, respectively. Four out of the eight genes from *N. glauca* (*g58903*, *g29887*, *g36929*, and *g72491*) cluster in a separate clade.

Some intolerant AA changes were also predicted to occur in this class, in three *N. glauca* genes in particular, which were generally highly expressed across all tissues. For example, *g6445* has a Ser substitution at position 132 (where all other sequences have Ala, *p*-value=9.625E-3), position 254 of *g36929* has been replaced by Ala (in place of a conserved Ser; *p*-value=4.16E-3) and *g29886* has a Met substitution at position 339 (where all remaining sequences have either Thr or Ser).

#### Alkane-forming pathway

##### CERl aldehyde decarbonylase component

Alkanes represent a major fraction of cuticular waxes, especially in *N. glauca*, where several long-chain hydrocarbons have been detected and can accumulate in considerable amount, up to 6 μg/mg FW [2], The protein CERl, CER3, and a cytochrome b5 component interact together to first reduce VLCFA-CoA to aldehydes and subsequently decarbonylate them to alkanes. There are seven genes annotated as CER1 in *N. glauca*, six in *N. sylvestris*, and seven in *N. tomentosiformis*. All CER1 genes from *N. glauca* are expressed across all tissues and leaf stages, with the exception of *g64050*, which is predominantly expressed in flowers and roots. Five (g18361, g30402, g50934, g58892, g64050) out of the seven genes in *N. glauca* show multiple AA changes, which are all predicted to impact protein function.

Additionally, we checked the aldehyde-generating component CER3, the cytochrome-b5 component, the cuticular lipid exporter PEC1, and the lipid transfer accessory factor LTPG; however, we found no interesting differences for these gene sequences.

### Analysis of genes of secondary metabolism

The three species of *Nicotiana* differ both in the total amounts and in the type of the predominant alkaloid accumulated. *N. sylvestris* has the highest content of total alkaloids in the leaves and accumulates nicotine primarily; on the other hand, *N. tomentosiformis* has a lower amount of total alkaloids (due to the deletion of the nic2 locus [42]) and accumulates nornicotine primarily; by contrast, *N. glauca* is one of the few *Nicotiana* species accumulating anabasine [43–45]. We therefore investigated the gene classes involved in the biosynthesis of pyridine alkaloids (nicotine, nornicotine, anabasine, etc.).

We focused on the class of diamino oxidases (DAO), which are generally involved in the oxidation of various amines, producing aminoaldehydes and hydrogen peroxide. Most of the DAOs are targeted to the apoplast, where they are involved in plant defense. Another group of DAOs, which are mainly targeted to the peroxisome, are active in polyamine catabolism, where they degrade putrescine to 4-aminobutanal. Some DAOs have been recruited in the synthesis of the pyrrolidine ring of nicotine. They can oxidize N-methylputrescine to 4-methylaminobutanal, the immediate precursor of the pyrrolinium cation. The genes encoding N-methylputrescine oxidases (MPO) evolved from DAOs acquiring new substrate specificity [15, 46]. We then checked if the amino acid polymorphisms and/or a particular pattern of tissue specificity of DAOs could represent factors explaining, at least in part, the low accumulation of nicotine in leaves of *N. glauca*. The *N. glauca*, *N. sylvestris*, and *N. tomentosiformis* genomes all contain three genes that are similar to the characterized *NtMPO1* and end with the amino acid sequences -SKL or -AKL with a peroxisomal localization [47, 48]. However, it is difficult to discriminate between canonical DAOs and MPOs based on sequence similarity without additional knowledge. In *N. tabacum*, for example, two DAO genes that share extremely high similarity (96%) perform different roles: one of them encodes a DAO involved in putrescine degradation (*NtDAO1*), while the other shows substrate preference for N-methylputrescine and is therefore a methylputrescine oxidase (*NtMPO1*) [46, 49]. On the basis of the phylogeny reported in Supplemental Figure 5, we can hypothesize that the clade including the bona fide *NtMPO1*, which shows substrate specificity for N-methylputrescine, contains homologous MPOs from the other species. The only gene from *N. glauca* in this clade, which could thus represent a bona fide MPO, as well, is *Ngla62825*. This was substantiated by checking six differing amino acids that distinguish MPOs from DAOs, which all supported the MPO function in this clade [46]. This *N. glauca* gene showed highest expression in the root [45].

### Variant analysis

To determine typical variance between the two *N. glauca* cultivars, *N. glauca* TW53 paired-end data representing approximately 30x coverage was mapped to the genome assembly, and we called variants using SAMtools with conservative settings. This resulted in the identification of 7,447,117 variants corresponding to a variant rate of approximately 0.2%.

The classification of the variants using SnpEff [50] revealed 3,833 variants that were rated as having a high severity (e.g., concerning stop codons and/or splice donor/acceptor sites). While a large proportion of these genes had no known function, it was noteworthy that the CER5 gene (*g6103*) was affected, as well. CER5 is an ABC transporter, which in *Arabidopsis* is responsible for wax export to the cuticle [51]. However, while this potential stop codon was very close to the C-terminus, it still fell in a region of high conservation.

### Field phenotyping experiments

We grew plants of *N. glauca* in a field trial in Madagascar in the 2015/2016 season and obtained between 4.7 g and 34 g of dried leaf material per plant. On average, this mirrors a harvest of about 22 g of leaf weight material per plant, representing between 188 kg/ha and 467 kg/ha of leaf yield. Compared with the 25 g of leaf material typically obtained from *N. tabacum*, this shows a generally lower yield, but we observed stark variation between different seed batches, suggesting that simple agronomic efforts might improve yield significantly. This is in line with a low germination capability of seeds, because these were not selected for high germination rates. However, despite the high content in secondary metabolites, we unexpectedly observed a need for fungicide and insecticide treatment, which may have caused this decrease in the yield of *N. glauca*. We also conducted a field trial in Dubai (UAE), where we grew *N. glauca* plants on dry soil using low irrigation volumes. In this case, the average leaf yield was approximately 450 g FW/plant, a result which is indicative of the potential of *N. glauca* to grow on marginal lands in conditions of partial drought.

## Discussion

Here we present the draft sequence of the tree tobacco *N. glauca*. Although the draft genome is still relatively fragmented, this could be addressed in the future using long reads to close sequencing gaps and to join smaller fragments. This could be achieved by the emerging nanopores [52] to yield a better representation of transposable elements and by using Hi-C and/or optimal mapping to assist in ordering scaffolds to chromosome scale units, as has been done to improve the *N. tabacum* genome [14], In any case, the large size of the *N. glauca* genome makes Illumina data-based assembly difficult, explaining the relatively low N50 obtained here. This draft genome allows exploring large regions of the genome unlike a previous assembly which as not annotated [53]. In addition, a BUSCO analysis and RNA-Seq mapping rates of 82-92%, together with the analysis of contained functional pathways, indicate that the genome is relatively complete in its gene space. Indeed, the BUSCO genome completeness analysis indicated that this draft genome is equivalent to other tobacco genomes. This makes the present genome an interesting resource that can be used (i) to find candidate genes involved in secondary metabolism and (ii) to better understand phylogenetic relationships in the *Nicotiana* genus.

The usefulness for secondary metabolism was highlighted when we analyzed a leaf developmental time course, including both metabolite and transcript data. We have found that terpenoids, which is a class of metabolites with a potential value, were generally upregulated during the developmental time course. As this trend was also visible from the gene expression analysis, it can be speculated that terpenoid synthesis is at least partially transcriptionally regulated in *N. glauca*, in line with modes of transcriptional regulation in other species [54, 55]. However, although one prime candidate for pathway analysis would be plant vitamin D synthesis, which could serve as a plant-derived source of vitamin D, we were not able to detect this metabolite in our *N. glauca* plants, rendering this approach infeasible at the moment. Possible reasons could be that *N. glauca* only produces this metabolite under stress conditions, such as stronger UV irradiation. This explanation would be consistent with the general induction of secondary metabolites by UV [56, 57] and plant-derived vitamin D in Solanaceous species in particular [31]. Another explanation might be that the usual occurrence of vitamin D is first observed indirectly by calcinosis in grazing animals [58] precluding proper plant identification. However, in the case of *N. glauca*, there is at least one reported case of direct identification of vitamin D from plant extracts [7].

The detailed analysis of leaf developmental time course from young to mature leaves - but non-senescing - revealed two clear, major trends: an upward and a downward trend in transcriptional regulation, as evidenced by clustering scaled data. Interestingly, while clustering time course data sets usually requires careful tuning of cluster numbers, in this case, two different measures clearly indicated two clusters to be optimal. This is in agreement with the partitioning of biological classes to the two clusters: the “downregulated” gene cluster mostly comprised cell proliferation-associated processes, such as cell cycle, chromatin, and cytoskeleton, confirming other studies on the model plant *Arabidopsis* [30]. This validates the current data set and underlines that the few classes identified as overrepresented in the “upregulated” gene clusters, such as terpenoid metabolism and SD-1 kinase, are likely under transcriptional regulation. Additionally, the compiled data set is thus likely to serve as a rich source to identify novel candidate genes in tobacco during leaf development, playing a particular role for biotechnological approaches. However, given high protein degradation dynamics during leaf development, at least in the model plant *Arabidopsis* [59], one needs to treat conclusions made only from transcripts with caution.

When analyzing cutin biosynthesis, the genome completeness was further underlined, as we generally found a similar number of genes for all investigated tobacco species. Somewhat surprisingly, we found relatively few non-conserved changes between *N. glauca* and other tobacco and Solanaceous species. One notable exception was CER1, a component of the aldehyde decarbonylase complex, which indicates that this protein might play a role in determining the composition of cuticular waxes in *N. glauca*, contributing to its partial drought resistance and to the accumulation of long-chain alkanes in its leaves.

Despite the considerable accumulation of alkanes in the leaves of *N. glauca*, its potential role as a biofuel crop deserves further investigation. When grown in field experiments under high irradiance, especially in Madagascar, the leaf yield was relatively low and characterized by a rather high degree of plasticity. When we compared our genomic data to a second independent cultivar, we were able to identify a relatively high percentage (0.2%, >7 million) of high-confidence nucleotide polymorphisms, suggesting that there is a relatively high degree of genetic diversity that could be exploited for genetic improvement. This value, although not as high as when compared with advanced draft genomes of *Solarium pennellii* tomatoes [17, 52], is on the high end if compared with the domesticated tomato, 5. *lycopersicum* [60].

## Potential implications

*N. glauca* represents a promising candidate species to serve as a source of feedstock for the biorefinery industry. It is a perennial species, able to grow on marginal lands in conditions of moderate drought and high temperatures; its leaves also may be harvested repeatedly during cultivation. Its potential for biofuel is based on the capacity of the plants to accumulate long-chain alkanes [2], The presented draft genome could thus become a resource to (i) further investigate the basis of the phenotypes that make *N. glauca* a potential biofuel crop, (ii) identify novel pathways of secondary metabolites, and (iii) to shed further light on phylogenomics in *Nicotiana*.

The exhaustive data set for a leaf developmental time course comprising both metabolites and transcripts could serve as a first entry point.

## Methods

### Plant material and field trials

*N. glauca* were obtained from Prof. Sandmann, Frankfurt (originally from the IPK seedbank), and the cultivar TW53 was available through the U.S. *Nicotiana* seedbank collection. They were grown in 30 cm diameter pots containing M2 professional growing medium under a 16-hour light/8-hour dark light regime in the greenhouse. For field trials, plants were grown in Madagascar.

### Genome sequencing and assembly

Short-insert “paired-end” libraries were prepared using the Illumina TruSeq DNA Sample Preparation Kit version 2 (Illumina, San Diego, CA). Long-insert “mate pair” libraries were prepared according to the Nextera Mate Pair Library Prep Kit (Illumina, San Diego, CA). All libraries were sequenced on an Illumina HiSeq-2500 using version 3 chemistry and flow cells with runs of 2 x 100 bases. Base calling and sample demultiplexing were performed using Illumina HiSeq Control Software and the CASAVA pipeline software.

Raw paired-end DNA reads were preprocessed with Trimmomatic (http://www.usadellab.org/cms/?page=trimmomatic) to remove sequencing adapters and low-quality reads from the 50 and 30 ends of the reads and to discard reads shorter than 50 bp. Raw mate-paired DNA reads were preprocessed with NxTrim (https://github.com/sequencing/NxTrim) to separate them into mate pairs and paired ends based on the presence of the Nextera adapter. The clean reads were then assembled into contigs using SOAPdenovo2 (http://soap.genomics.org.cn/soapdenovo.html) [18] with a k-mer of 75 and scaffolded by increasing library size. Gaps resulting from the scaffolding were closed using GapCloser (http://soap.genomics.org.cn/soapdenovo.html), and all sequences shorter than 200 bases were discarded from the final assemblies. Remaining gaps with overlapping flanking regions were then closed using in-house scripts. After closing the gaps, singletons were used as queries in a BLAST-based algorithm against the scaffolds. They were eliminated if the match level was greater than 99% to avoid artificial duplications of short sequences.

### Genome annotation

To this aim, we trained AUGUSTUS by public EST and RNA-Seq data and the 26 new RNA sequencing samples generated in this project covering a variety of developmental leaf stages and tissues.

For a functional gene annotation and assignment to MapMan classes, we used the Mercator version X4.03 pipeline. To enable cross-species comparison, we used the same pipeline version to annotate other Solanaceous species, as well, using the same online pipeline. The only difference was that for *N. glauca*, we used a six-frame translation of nucleotide sequences to improve sensitivity for partial gene models. We visualized data using a custom-written D3 application derived from a sunburst visualization from web visits (https://usadellab.github.io/dataviz/sunburst/index.html). We further investigated MapMan classes manually that showed significant differences in their proportion in *N. glauca* in comparison with the other species.

### SNP calling

For SNP calling, we trimmed raw read data using Trimmomatic [61] in paired-end mode using default settings. Then we aligned the trimmed reads using Bowtie2 and preprocessed them using Picard SortSam. We called variants using the SAMtools mpileup script. We determined high-quality variants using vcfutils, applying a quality filter of 30 or above and requiring a coverage (“DP“) of 6 or above. Finally, we annotated variants with SnpEff [50].

### Repetitive elements

We detected and classified transposons by a homology search against the REdat_9.5_Eudicot section of the PGSB transposon library [62], We used the program vmatch (http://www.vmatch.de), a fast and efficient matching tool, with the following parameters: identity>=70%, minimal hit length 75 bp, seed length 12 bp (exact command line: -d -p -l 75 -identity 70 -seedlength 12 -exdrop 5). We filtered the vmatch output for redundant hits via a priority-based approach, which assigns higher scoring matches first and either shortens (<90% coverage and >=50bp rest length) or removes lower-scoring overlaps to obtain an overlap free annotation.

### RNA sequencing

*N. glauca* plants were grown in the greenhouse, and roots, stems, and flowers were harvested in duplicates. In addition, leaves were harvested at 10 developmental stages in duplicates. We extracted RNA using TRIzol and a RNA extraction kit (Qiagen), as previously described [19], and paired-end sequencing libraries were sequenced in the Illumina Core Center at the Max Planck Institute in Cologne using standard procedures. For differential expression analysis, we mapped reads to the reconstructed transcriptome using salmon [22], Subsequently, we analyzed data using edgeR [27] using RobiNA [26].

### Analysis of selected gene classes

All predicted protein-encoding genes from *N. glauca*, *N. sylvestris*, and *N. tomentosiformis* were submitted to Mercator 4 v.0.2 to get a functional annotation based on MapMan bins. The process is based on the assignment of a gene functional category to the query sequences based on a similarity search to a number of reference databases [23]. The results from Mercator allowed us also to compare the different number of genes assigned to each bin category for the three species. For each Mercator gene class, the longest isoforms were selected from *N. glauca*, *N. tomentosiformis*, and *N. sylvestris* to build multialignments and phylogenetic trees using the online tool at http://phvlogenv.lirmm.fr [63]. For each gene class, the full protein alignment along with its phylogenetic tree were also used as inputs for MAPP [32] in order to get a prediction of the impact of missense mutations on protein function. The approach is based on the amount of physicochemical variation in each position of a multialignment built from a set of homologous sequences.

### Metabolite profiling

We extracted stem and leaf material of *N. glauca* in ball-mill, pre-injection, injection, and analyses steps for gas chromatography coupled to mass spectrometry, and polar and apolar liquid chromatography coupled to mass spectrometry were carried out by following the protocols previously developed [64–66] as well as [67] for isoprenoids exactly. We annotated peaks by reference to database entries’ [68] previously annotated peaks in tobacco tissues [69] and reported metabolites in accordance with suggested standards [70]. Detailed steps for metabolic analysis have been detailed in [19].

### Accession codes and cDNA availability

Genome sequence data for *N. glauca* have been deposited in the DDBJ/EMBL/GenBank nucleotide core database under the accession code MDKF00000000. Genome and transcriptome sequencing data for *N. glauca* have been deposited in the GenBank Sequence Read Archive under BioProject PRJNA335307.

## Acknowledgments

The research from the MultiBioPro project leading to these results has received funding from the European Community’s Seventh Framework Programme for research, technological development, and demonstration under grant agreement 311804. BU wants to thank the Horizon2020 program GoodBerry under grant agreement 679303, which supported tool development for functional analysis. AK and PDF thank the Biotechnology and Biological Sciences Research Council for the Integrated Biorefining Research and Technology Club programme. BU, HG, MS, and KM thank the Bundesministeriums fur Bildung und Forschung program deNBI 031A536. SP was supported by an Australian Research Council Future Fellowship grant (FT160100218) and a University of Melbourne International Research and Research Training Fund grant. Finally we thank Marie Bolger, Rainer Schwacke, and Mario Stanke for supplying data and materials.

